# A model of actin-driven endocytosis explains differences of endocytic motility in budding and fission yeast

**DOI:** 10.1101/2021.07.20.453152

**Authors:** Masoud Nickaeen, Julien Berro, Thomas D. Pollard, Boris M. Slepchenko

## Abstract

A comparative study (Sun et al., eLife, 2019) showed that the abundance of proteins at sites of endocytosis in fission and budding yeast is more similar in the two species than previously thought, yet membrane invaginations in fission yeast elongate two-fold faster and are nearly twice as long as in budding yeast. Here we use a three-dimensional model of a motile endocytic invagination (Nickaeen et al., MBoC, 2019) to investigate factors affecting elongation of the invaginations. We found that differences in turgor pressure in the two yeast species can largely explain the paradoxical differences observed experimentally in endocytic motility.

## Introduction

As in plant cells, endocytosis in yeast cells occurs under high turgor pressure estimated to be ~10 atm in fission yeast (Basu et al., 2014; Lacy et al., 2018), and about a fifth to half of that in budding yeast (de Maranon et al., 1996; Schaber et al., 2010). Transient assembly of small, dense networks of actin filaments at endocytic sites (termed ‘actin patches’) is necessary for robust endocytosis in yeast ((Aghamohammadzadeh and Ayscough, 2009; Basu et al., 2014)), suggesting that assembly of an actin patch around a nascent invagination of the plasma membrane may generate a pulling force sufficient to elongate the invagination under such pressures. Many studies have investigated the mechanisms of force generation at endocytic sites in yeast (Carlsson and Bayly, 2014; Scher-Zagier and Carlsson, 2016; Carlsson, 2018; Lacy et al., 2018; Kaksonen and Roux, 2018; Mund et al., 2018; Nickaeen et al., 2019).

A comparative study of endocytosis in fission and budding yeast (Sun et al., 2019) found that with the exception of two-fold more polymerized actin in fission yeast the abundances of proteins participating in patch assembly are more similar than previously thought. Nevertheless, elongation of an endocytic invagination is two-fold faster (~52 nm/s) in fission yeast than in budding yeast (~ 24 nm/s). The fast elongation rates in both yeasts indicate that driving forces generated at endocytic sites substantially overpower resistance from turgor pressure, not just withstand it. For the turgor pressure of 10 atm, a quick estimation yields a resisting force of ~3,000 pN acting on a cylindrical tubule with a typical radius of ~30 nm (Carlsson, 2018; Lacy et al., 2018).

Our simulations of a spatial model of a motile invagination estimated that an actin filament network around a cylindrical tubule with a radius of 30 nm can generate tangential pulling forces of ~ 2500 pN (Nickaeen et al. 2019). With such forces, the tubule can withstand a turgor pressure of ~ 9 atm and elongate slowly below this threshold, for example, at only ~ 2 nm/s against a turgor pressure of ~7 atm. Yet the model made another prediction that the assembling patch would also generate normal forces, which squeeze the tubule at its base and stretch it at its middle, thus transforming the invagination shape from cylindrical to a flask-like (or ‘head-neck’) as observed in electron micrographs of budding yeast (Kukulski et al., 2012; Buser and Drubin, 2013). Previous modeling studies also predicted endocytic invaginations with head-neck shapes (Dmitrieff and Nédélec, 2015; Ma and Berro, 2021). Since the resisting force is the product of turgor pressure and the cross-sectional area at the base of the invagination, the transition to the head-neck shape may dramatically reduce resistance due to turgor pressure, leading to faster elongation rates even for turgor pressures of ~10 atm.

Our previous study, as work of others (Carlsson and Bayly 2014; Mund et al. 2018), modeled the invaginations as spherocylinders. In those simulations, we approximated the reduction in resistance due to the putative shape change by replacing fixed resistance with a resistance decreasing over time. The elongation rate increased four-fold, to ~8 nm/s, which is still significantly lower than the 25-50 nm/s range reported by Sun et al.

In this study, we solved our model in geometries mimicking flask-like invaginations. The solutions yielded higher driving forces, because, in contrast with cylindrical shapes, for which pulling forces are viscous in nature, active stresses also contribute to the forces driving flask-like invaginations. Consistent with the experimental data (Sun et al., 2019), invagination is faster and deeper at high turgor pressures, because at higher initial resistance, more actin filaments accumulate by the time the driving force overcomes the resistance, producing higher driving forces during elongation.

## Model and Methods

Our model, described in detail in (Nickaeen et al., 2019), combines kinetics of actin nucleation, polymerization, and turnover constrained by counts of each participating protein over time (Berro et al., 2010), and the mechanics of the assembling filamentous meshwork approximated as that of a visco-active gel (Kruse et al. 2005; Prost et al., 2015). Mathematically, the model consists of advection-reaction equations governing densities of proteins involved in patch assembly (see Eq (S1) in *Supplemental Text*), and a force-balance equation yielding actin velocities (Eq (S2) of *Supplemental Text*). In the advection-reaction equations, reaction rates and rate constants from the experimental literature (Berro et al., 2010; see also Table S1 *in Supplemental Text*) were modified to reflect effects of forces and filament densities on polymerization kinetics. In the force-balance equation, active repulsive stresses caused by impingement of polymerized subunits on existing filaments are proportional to the square of local density of polymerized subunits, and viscous stresses due to entanglement and crosslinking of branched filaments are assumed to be largely proportional to the square of local density of polymerized subunits and to the local average length of filaments (Gardel et al., 2003; Nickaeen et al., 2019; see also Eq (S3) in *Supplemental Text*). The coefficients determining scales of the stresses were inferred from rheological properties of actin filament networks (Gardel et al., 2003; Kasza et al., 2010; Tseng and Wirtz, 2004; Mullins et al., 1998).

Through interactions with the membrane, the flow of polymerized actin exerts on the invagination driving forces parallel to its axis. The resultant driving force (*f*_drive_) and resistance due to turgor pressure (*f*_resist_) are the factors determining elongation rate in our model.

Solving for the invagination shape would involve membrane mechanics and require reliable information about composition and rheological characteristics of both the membrane and its protein coat. Instead, we solve the model for head-neck invaginations whose head and neck radii do not change as the neck elongates, on the assumption that all points of the membrane move with the same speed *u*(*t*) described by a linear force-velocity relation,

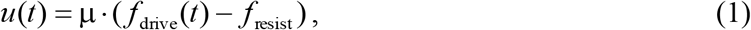

where *μ* is the mobility coefficient of the invagination. For the invagination to elongate, the driving force must exceed the resistance of turgor pressure, therefore Eq (1) holds only for *f*_drive_(*t*) ≥ *f*_resist_, whereas for *f*_drive_(*t*) <*f*_resist_, *u*(*t*) = 0.

Nucleation-promoting factors (NPFs) – the proteins that stimulate Arp2/3 complex to nucleate branched actin filaments - reside on the invaginating membrane, where they concentrate in narrow rings around the membrane. Fission yeast has two rings of NPFs, one that remains in the initial position at the base of the invagination at the plasma membrane, while the other moves with the tip of the tubule (Arasada and Pollard, 2011; Arasada et al., 2018), whereas budding yeast has one ring of NPFs that remains near the base of the invagination (Mund et al., 2018). As with cylindrical shapes (Nickaeen et al., 2019), simulations of both arrangements of NPFs on head-neck surfaces yielded similar elongation rates and tip displacements. To avoid duplication, we employ the two-zone arrangement of NPFs characteristic of fission yeast to illustrate our findings throughout this paper.

The coupled system of the force-balance equation, reaction-transport equations, and Eq (1) was solved using a moving-mesh solver of COMSOL, a software package for solving spatial multiphysics problems on finite element meshes (COMSOL Multiphysics, 2015). Because the invaginations were modeled as axially symmetric, computations were simplified by reducing the original three-dimensional (3D) problem to an equivalent two-dimensional model formulated in (*r, z*) coordinates. The supplemental materials of our previous paper (Nickaeen et al., 2019) provide more details about the model and its numerical solutions.

## Results

### 1. Changing the shape of the invagination from cylindrical to flask-like amplifies driving forces, yielding faster elongation and longer displacements

Our previous study revealed that an assembling actin patch not only produces a pulling force but also generates orthogonal forces, which squeeze the base of the invagination and stretch its middle (Figure 1A). As actin filaments accumulate around an initial, cylindrical invagination, squeezing forces at the base of the invagination reach the magnitude amounting to additional pressure of ~ 8.5 atm (Figure 1B). It is therefore likely that at some point during patch assembly, the nascent spherocylindrical invaginations, such as the one shown in Figure 1A, acquire head-neck shapes (Figure 1C). Such shapes were indeed observed in electron micrographs of invaginations in budding yeast (Kukulski et al., 2012; Buser and Drubin, 2013). Flask-like shapes were also predicted theoretically for a membrane elongated under high turgor pressure by a point force exerted at its tip (Dmitrieff and Nédélec, 2015; Ma and Berro, 2021).

**Figure 1.**
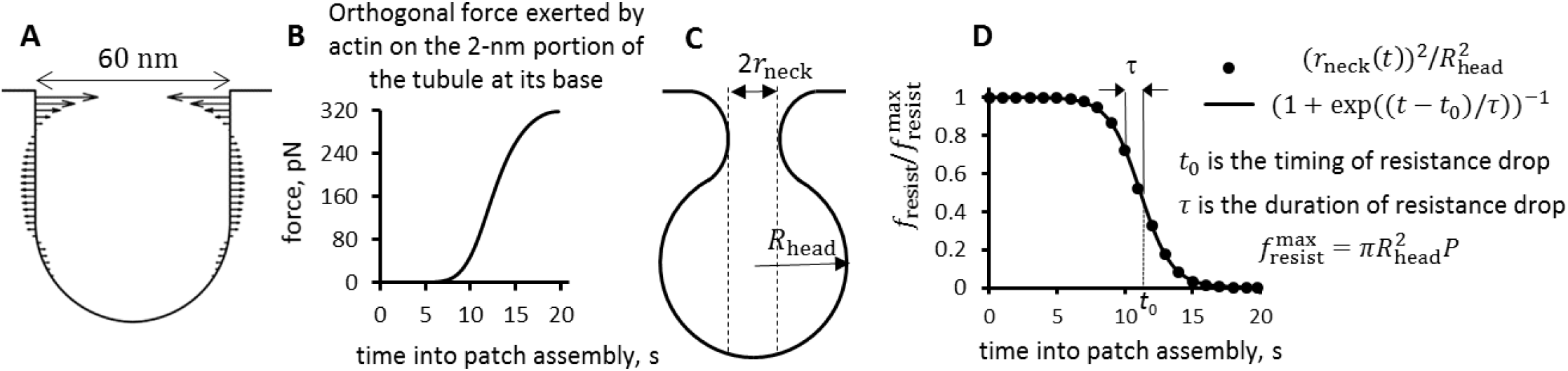
An assembling endocytic actin patch exerts strong squeezing forces at its base of an initial invagination. (A) Distribution of orthogonal forces shown for an (*r, z*)-section of a nascent invagination, adapted from simulation results in (Fig.5A of Nickaeen et al., 2019). (B) Time dependence of squeezing force applied to the 2 nm wide portion at the base of the nascent tubule in (A), from a simulation described in Fig.5A of (Nickaeen et al., 2019). The force of 320 pN applied to a 2 nm wide band with the radius of 30 nm amounts to additional pressure of ≈8.5 atm. (C) Cartoon of a head-neck (flask-like) shape with radii *r*_neck_ and *R*_head_; since turgor pressure *P* is isotropic, *f*_resist_ for this shape diminishes to 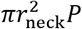. (D) Graph of *f*_resist_(*t*) normalized to 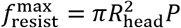 (dots) is well approximated by function (1 + exp((*t* – *t*_0_)/*τ*))^−1^ (solid curve) with the fitted timing (*t*_0_) and duration (*τ*) of the resistance decrease caused by shape change.

Because turgor pressure is isotropic and thus pushes on the upper surface of the invagination head as well, the transition from the cylindrical shape to the flask-like one, such as in Figure 1C, *f*_resist_ in Eq (1) to *f*_resist_(*t*) =(π(*r*_neck_*t*))^2^*P*, where *P* is the turgor pressure and *r*_neck_(*t*) is the time-dependent neck radius. Normalizing *f*_resist_(*t*) to its maximum, 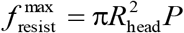, yields 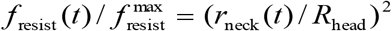, where *R*_head_ is the head radius. The dots in Figure 1D represent normalized resistance for the *r*_neck_(*t*) derived from the orthogonal forces in Figure 1B, assuming Hookean elasticity of the protein coat (we also assumed for simplicity that as the force approaches its maximum, *r*_neck_ → 0). Notably, a function (1 +exp((*t* – *t*_0_)/τ)) ^−1^ with the suitable timing *t*_0_ and duration τ of the resistance descent (solid curve in Figure 1D), provides a close fit. Note that, because the reduction of resistance occurs due to shape change, *t*_0_ also represents the time around which the transition to a flask shape takes place. Based on these observations, we approximate *f*_resist_ by the following function of time,

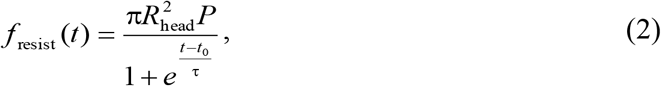

and treat *t*_0_ and τ as model parameters (see also Eq (2*) in the next subsection).

However, simulated cylindrical invaginations did not elongate as fast as observed experimentally, if only the reduction of resistance is taken into account (see *Introduction*). Seeking faster elongations, we noted that the shape change should also affect the driving force. Indeed, driving forces originate from stresses in the actin filament meshwork surrounding the invagination, and while the forces pulling cylindrical invaginations derive from viscous stresses, the driving forces exerted on curved invaginations would have additional contributions from active stresses due to the actin flow pushing on the upper surface of the invagination head. This would not only increase the driving force during elongation, but might also make elongation rates more sensitive to the invagination mobility coefficient μ (Eq (1)).

The elongation rates of cylindrical invaginations are essentially insensitive to μ, because the viscous driving forces depend on shear rates of actin flow at the invagination that are lower for invaginations with higher mobility. As a result, the increase of μ is counterbalanced by the drop in the viscous pulling force, leaving the elongation rate *u*(*t*) virtually unchanged (Nickaeen et al., 2019). In contrast, the forces driving a flask-like invagination might be less sensitive to reductions in shear rates. The reason is that active stresses are by definition independent of shear rates, and so are their contributions to the driving force.

We tested these hypotheses by solving the model in geomteries mimicking invaginations with head-neck shapes. To maximize elongation rates, we explored shapes with small ratios *r*_neck_/*R*_head_ (the head radius was fixed at 30 nm). Given the reported similarity of protein abundances in fission and budding yeast, we ran all simulations with a same set of reaction rate constants and abundances of soluble proteins taken from (Berro et al., 2010; Nickaeen et al., 2019). Similarly, we uniformly applied coefficients setting the scales for active and viscous stresses as they were derived in our previous study. Results were consistent with our expectations, as exemplified by a solution obtained for an invagination with *r*_neck_/*R*_head_ = 0.1 (Figure 2), using the same parameters as for the cylindrical invagination in our previous study (Fig. 7 in ref), with the exception of the mobility coefficient value μ = 0.08 nm/(s·pN), which is twice the one used for the cylindrical invagination^1^.

**Figure 2.**
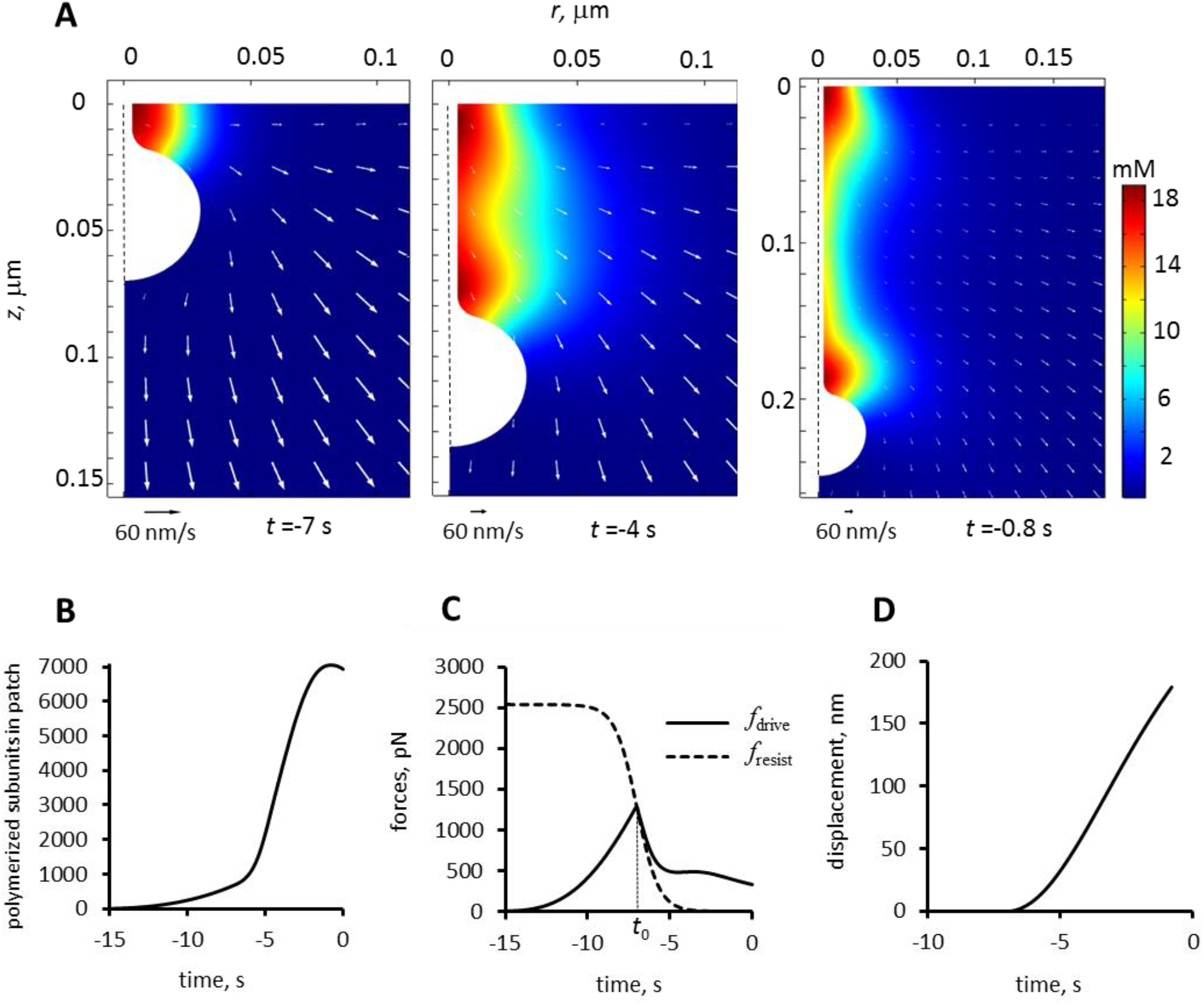
Modeling an elongating invagination with head-neck shape. We simulated elongation of the head-neck invagination described by r_neck_/ *R*_head_ = 0.1, using the following parameters: *P* = 9 atm, *t*_0_ = −7 s (13 s into patch assembly), *τ* = 0.66 s, and *μ* = 0.08 nm/s/pN. Similar to experimental studies, the time here and below is offset by −20 s, so that the times of actin peaks would be around time zero. (A) The (*r, z*) sections of 3D distributions of actin densities (pseudo-colors) and velocities (white arrows) shown for three times: (left) beginning of elongation; (middle) actin half maximum, and (right) actin peak. Extracellular space is white. Velocity scale bars correspond to 60 nm/s. (B) Polymerized actin as a function of time. The numbers of subunits were counted inside a growing cylinder embedding the moving invagination: *r_cyl_* = 0.06 μm, 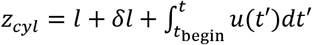, where *t*_begin_ denotes beginning of elongation and lengths are in μm: the initial invagination length *l* = 0.07 μm and short extra length *δl* = 0.003 μm. (C) Time dependence of driving force (solid curve) and resistance due to turgor pressure described by Eq (2) (dashed curve); the invagination mobility used is *μ* = 0.08 μm/s/pN. (D) Displacement is ~ 180 nm at the time actin peaks, which is thought to coincide with scission, yielding average elongation rate ~ 30 nm/s, over the duration of elongation, *t*_scission_ – *t*_begin_.

The snapshots of the (*r*,*z*) sections in Figure 2A correspond to beginning of elongation (left), actin half-maximum (middle), and actin peak (right) (note that we used different spatial scales in Figure 2A, right, to accommodate the longer displacement). The time in this figure and below are offset by −20 s, so that on the transformed scale, actin peaks, which are thought to coincide with the scission of endocytic vesicle (Sun et al., 2019), occur around time zero (Figure 2B). Thus, the offset readings are close to times before scission.

A nascent invagination begins to elongate (Figure 2D) at the time *t*_begin_ when the solid curve in Figure 2C depicting the driving force intersects with the dashed graph of the resistance of turgor pressure. In this particular example, *t*_begin_ coincides with *t*_0_, but generally may deviate from it, as discussed in the next subsection. The driving force, rising at *t* < *t*_begin_, starts ‘shadowing’ the descending *f*_resist_(*t*) immediately after *t*_begin_ (Figure 2C). This transition is largely a result of an abrupt drop of the viscous component of *f*_drive_ caused by the decrease in the shear rate, as the moving invagination catches up with the surrounding actin flow.

Assuming the vesicle scission coincides with the time of maximum actin, the simulated duration of elongation is 6.2 s. Notably, the maximum displacement 179 nm (Figure 2D) yielded by the model is within the range of displacements measured in yeast: ~125 nm in budding yeast, and ~200 nm in fission yeast (Sun et al., 2019). The average elongation rate is then 179 nm/6.2 s = 28.9 nm/s. The maximum elongation rate, achieved at *t* = −3.3 s, is 38 nm/s. Both the average and maximum values are in the range of speeds reported by Sun et al. Thus, solving the model in geometries mimicking curved flask-like invaginations and with the amplified invagination mobility produced elongation rates and displacements comparable with experimental data.

The factors contributing to the greater speeds and displacements can be elucidated further by comparing simulations that differ only by invagination shape and mobility. For this, we reran the simulation described above with μ = 0.04 nm/(s·pN) and compared results of both simulations with the solution of the same model for the cylindrical invagination with μ = 0.04 nm/(s·pN) from our previous study (Figure 3).

**Figure 3.**
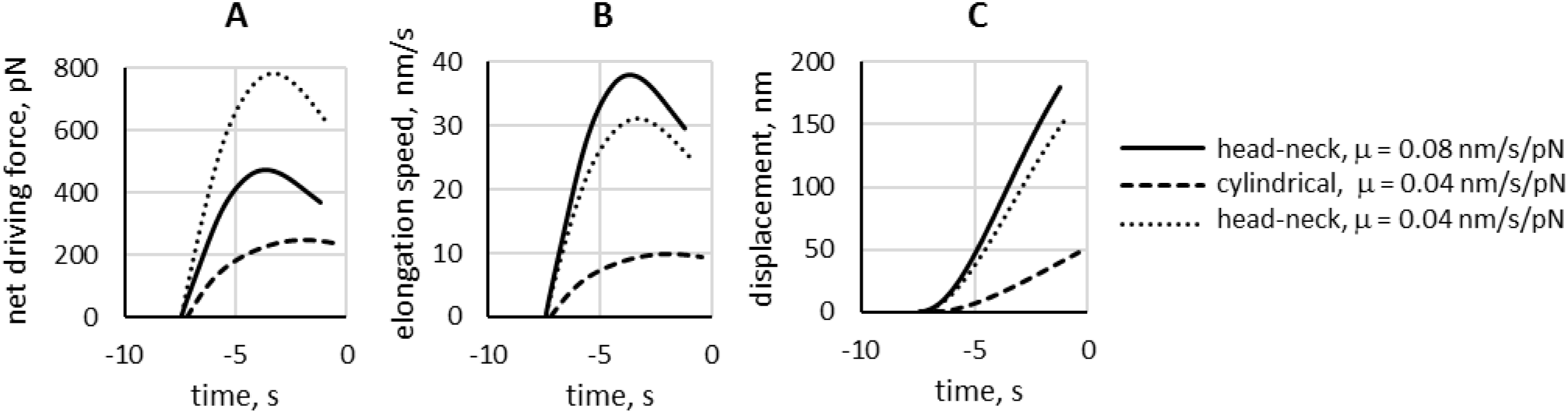
Invaginations with ‘head-neck’ shape have higher driving forces than cylindrical invaginations, yielding faster elongation and longer displacements. Dependence of (A) net forces, *f*_net_ = *f*_drive_ – *f*_resist_, (B) elongation rates and (C) displacements on the invagination geometry and *μ*. The model was solved with the same parameters (with the exception of invagination mobility) for the invagination with head-neck shape (*r*_neck_ /*r*_head_ = 0.1) and mobility coefficients *μ* = 0.08 μm/s/pN (solid curves) and *μ* = 0.04 μm/s/pN (dotted curves) and for the cylindrical invagination with *μ* = 0.04 μm/s/pN (dashed curves). The head-neck invagination began to move at *t*_begin_ = −7 s, and the cylindrical invagination started to elongate at *t*_begin_ = −6.8 s. Vesicle scission, terminating elongation, is thought to occur when polymerized actin reaches its maximum. Peak numbers of actin molecules for the head-neck invagination were achieved at *t*_scission_ = −0.8 s for *μ* = 0.08 μm/s/pN and − 0.6 s for *μ* = 0.04 μm/s/pN, and for the cylindrical invagination with *μ* = 0.04 μm/s/pN) - at *t*_scission_ = 0. The displacements at *t*_scission_ were 179 nm (solid curve), 153 nm (dotted curve), and 42 nm (dashed curve).

Changing the shape from cylindrical to flask-like produced a three-fold higher maximum of the net driving force, *f*_net_(*t*) = *f*_drive_(*t*) – *f*_resist_(*t*) (dashed and dotted curves in Figure 3A). Because all three solutions were obtained with the same *f*_resist_(*t*), the higher *f*_net_ reflects the anticipated increase of *f*_drive_. Note that simulations of the head-neck invagination with the higher mobility of 0.08 μm/s/pN result in lower *f*_net_ (dotted and solid curves in Figure 3A), because the contributions of viscous stresses to *f*_drive_ depend on shear rates, which are lower for higher μ, as discussed above. Yet overall, the higher μ produced noticeably faster elongation rates *u*(*t*) = μ · *f*_net_(*t*) (solid and dotted curves in Figure 3B), which validates the prediction that the elongation of curved invaginations is more sensitive to changes of *μ* than that of the cylindrical invaginations (Nickaeen et al., 2019). The maximum elongation rates in Figure 3B were as follows: (solid curve) 38.0 nm/s, (dotted curve) 31.2 nm/s, and (dashed curve) 9.9 nm/s. The higher elongation rates translate into larger maximal dispacements 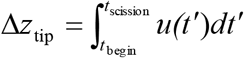 plotted in Figure 3C, where *t*_scission_ is intepreted as the time of the peak of actin. The legend of Figure 3 gives the values of *t*_begin_ and *t*_scission_ for each of the solutions. Based on these data, the average elongation rates 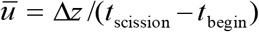 are (solid curve) 28.9 nm/s, (dotted curve) 23.9 nm/s, and (dashed curve) 7.7 nm/s.

In this study, we do not explicitly model the transition of a nascent cylindrical invagination into a flask shape. Instead, initial invaginations in our simulations are aready of a head-neck shape, in obvious contradiction with Eq (2), which assumes that the shape transformation occurs around time *t*. However, the errors caused by this inconsistency are likely to be small. Indeed, according to Figure 3, the cylindrical and head-neck invaginations, simulated with the same *f*_resist_(*t*), begin to elongate at approximately same time, suggesting that during the time before the elongation, *f*_drive_(*t*) is similar for all shapes. Simulations of invaginations with different *r*_neck_ /*R*_head_ and *f*_resist_(*t*), described in the next subsection, also show that the pre-elongation *f*_dnv_ e(*t*) are virtually independent of shape (see Figure 4A in the next section).

**Figure 4.**
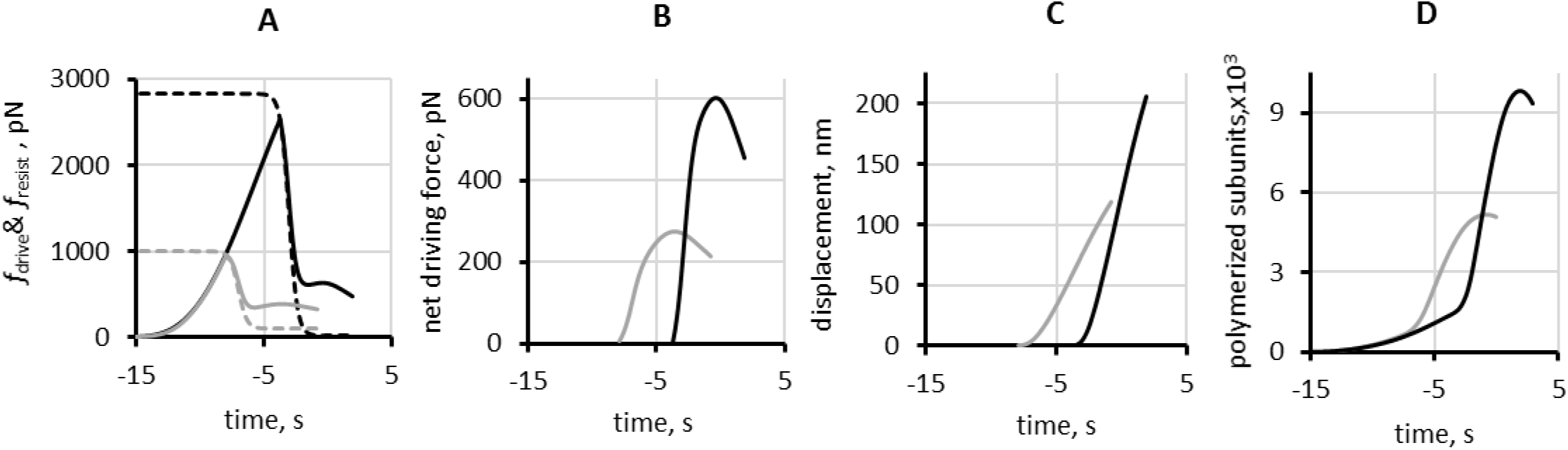
Output of simulations of models using optimized parameters for budding yeast (grey curves) and fission yeast (black curves).

(A) Solid curves are *f*_drive_(*t*), dashed curves are graphs of *f*_resist_(*t*, see Eq (2*); (B) net forces *f*_net_(*t*) = *f*_drive_(*t*) – *f*_resist_(*t*), with *f*_drive_(*t*) and *f*_resist_(*t*) from (A); (C) displacements; (D) polymerized actin, with the maxima of 9826 subunits in fission yeast and 5165 subunits in budding yeast.

The budding yeast results are from the simulation with *r*_neck_/*R*_head_ = 1/3. Values of *t*_0_ used in the simulations (Table 3) are ‘near-saturation’, as evident from graphs in (A). Simulated displacements (C) and elongations rates (Table 4) are in reasonable quantitative agreement with experimental data of Sun et al. (Sun et al., 2019). Results for polymerized actin in (D) predict that at scission, actin patches in budding yeast have fewer polymerized actin subunits than in fission yeast (see discussion in next subsection).

This gives us leeway in defining shapes during the time before the elongation. By using at *t* < *t*_begin_ a fixed head-neck shape that would arise from shape transformation, we capture a makeup of the driving force in terms of active and viscous components at *t* = *t*_begin_. Unlike the total driving force, the pre-elongation active and viscous components depend on shape significantly, and the composition of *f*_drive_(*t*) at *t* = *t*_begin_ in terms of these components defines the behavior of the driving force during elongation and, ultimately, the characteristics of motility of the invagination.

### 2. The head-neck invaginations elongate faster and to greater depths under higher turgor pressures

One might assume that elongation would be fastest if the resistive force were fixed at a low value. However, simulations of a head-neck invagination moving against a constant resistive force of 28.3 pN yielded average elongation rates of only ~ 15 nm/s (Table 1) in spite of resistance two orders of magnitude lower than experienced by a cylindrical invagination under a turgor pressure of 10 atm.

**Table 1.** Average rates of elongation of a head-neck invagination (*r*_neck_ /*R*_head_ = 0.1; *R*_head_ = 30 nm) against resistive force of 28.3 pN with three invagination mobility coefficients and different arrangements of NPFs.

| Mobility coefficient, nm/s/pN | Average elongation rate, nm/s |  |  |
| --- | --- | --- | --- |
|  | Two rings of NPFs: one fixed at base, the other moving with invagination | One ring of NPFs fixed on invagination at base | One ring of NPFs on flat cell membrane surrounding the invagination opening (as in Mund et al., 2018) |
| 0.01 | 8.1 | 7.3 | - |
| 0.04 | 12.6 | 13.6 | - |
| 0.08 | 14.3 | 16.6 | 13.2 |

This counterintuitive behavior arises, because the driving force overcomes the resistance early during patch formation. At this point, the actin flow is slow and the viscous component dominates the driving force, resulting in slow elongation. When the active component subsequently grows stronger, the viscous component decreases and even becomes resistive.

These factors keep the overall driving force down, yielding relatively slow rates during the entire elongation. Conversely, a stronger, and longer lasting initial resistance gives the driving force a chance to build up and may produce faster elongation rates and deeper invaginations, once the driving force overcomes the resistance.

To test this prediction, we compared simulations with lower turgor pressure representative of budding yeast and the other with high turgor pressures characteristic of fission yeast. We also varied the invagination mobility coefficient μ and the parameter *t*_0_ approximating the timing of shape change; as above, *t*_0_ was offset by −20 s. In all simulations, the neck-head ratio was *r*_neck_ / *R*_head_ = 0.1 and the duration of shape change was τ = 0.33 s. Table 2 summarizes the simulation parameters and results, including: the duration of elongation computed as *t*_scission_ – *t*_begin_ with *t*_scission_ intepreted as the time of the peak of actin, the maximum displacement defined as the displacements at *t* = *t*_scission_, the average elongation rate (the ratio of the maximum displacement and the corresponding duration of elongation), and the peak amount of polymerized actin computed as described in the legend of Figure 2.

**Table 2.** Parameters and results of simulations with varying *P*, *t*_0_, and μ.

|  | Simulations | Simulation parameters |  |  | Simulation results |  |  |  |  |
| --- | --- | --- | --- | --- | --- | --- | --- | --- | --- |
| | | Turgor pressure ($P$), atm | Timing of resistance descent ($t_0$), s | Mobility coefficient ($\mu$), nm/(s·pN) | Duration of elongation, s | Maximum displacement, nm | Average elongation rate, nm/s | Maximum elongation rate, nm/s | Maximum number of patch actin subunits, $\times 10^3$ |
| Lower turgor pressure | 1 | 5 | -6 | 0.08 | 5.8 | 185 | 32.0 | 39.3 | 7.3 |
|  | 2 | 3.5 | -7 | 0.08 | 6.2 | 176 | 28.4 | 34.8 | 6.2 |
|  | 3 | 3.5 | -8.5 | 0.08 | 7.5 | 185 | 24.6 | 29.7 | 5.2 |
|  | 4 | 3.5 | -8.5 | 0.16 | 7.1 | 197 | 27.8 | 33.4 | 5.4 |
|  | 5 | 3.5 | -8.5 | 0.32 | 6.7 | 200 | 29.8 | 35.8 | 5.5 |
| Higher turgor pressure | 6 | 9 | -4 | 0.08 | 5.2 | 202 | 38.8 | 46.8 | 9.4 |
|  | 7 | 9 | -4 | 0.16 | 5.1 | 222 | 43.5 | 52.9 | 10.2 |
|  | 8 | 9 | -4 | 0.32 | 5.1 | 236 | 46.8 | 56.7 | 10.6 |
|  | 9 | 10 | -3 | 0.32 | 5.0 | 237 | 47.7 | 59.0 | 11.2 |

As predicted, the simulated elongation rates were faster and the displacements longer at higher turgor pressures. Also, the timing of the resistance reduction *t*_0_ correlated positively with elongation rates (see simulations 2 and 3) and the simulation outcomes were only modestly sensitive to μ, consistent with results in Figure 3B and C.

While both *P* and *t*_0_ influence the motility of invaginations (Table 2), the impact of *P* is overarching, because turgor pressure limits the effect of *t*_0_ on both *t*_begin_ and *f*_drive_(*t*) at *t* > *t*_begin_. Indeed, both *f*_drive_(*t*_begin_) and *t*_begm_ saturate with increasing *t*_0_ at the values controlled by turgor pressure, 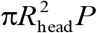 and 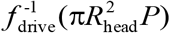, respectively (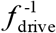 stands for the inverse function of *f*_drive_(*t*)).

This is because the time *t*_begin_ is the solution of *f*_drive_(*t*) = *f*_resist_(*t*) and 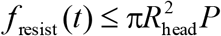 (Eq (2)); note also that *f*_drive_(*t*_begin_) is the absolute maximum of *f*_drive_(*t*) (Figures 1C,D and 3A,C). The saturation begins at 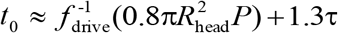, for which 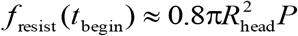 (see *Supplemental Text*). Interestingly, the ‘near-saturation’ values of *t*_0_ used in simulations 1, 2, and 6-9 (Table 2) resulted in longer elongations at lower turgor pressures, which is consistent with results in Fig. 3 of (Sun et al., 2019).

The simulation results in Table 2 agree qualitatively with the observations that invaginations in budding yeast move slower and are shorter than in fission yeast (Sun et al., 2019), which is consistent with the different turgor pressures explaning these differences in the two species. Quantitatively, though, our simulations significantly overestimated the displacements in budding yeast. Note also that the elongation rates in simulations 3 and 9 are similar to those in budding and fission yeast, but they were obtained with the four-fold difference of the invagination mobility in the two yeasts, which may not be realistic.

We ran additional simulations with varying model parameters to find better agreement with reported data. Simulations with parameter sets termed ‘optimized’ (Table 3) yielded results more consistent with the measurements by Sun et al. (Table 4). In these sets, mobility coefficients μ and durations of resistance reduction τ are similar for both yeasts. Parameters that differ include turgor pressure *P*, pressure-dependent ‘near-saturation’ values of *t*_0_, and neck-head ratios *t*_neck_/*R*_head_ (recall that all simulations in Table 2 were run with the same *r*_neck_/*R*_head_ = 0.1). The shapes of the invaginations may differ in the two species because of different normal forces or rigidities of the protein coat, or both. The coat rigidities are unknown, but the compressing force might be weaker in budding yeast, since turgor pressure, that contributes to squeezing the

**Table 3.**
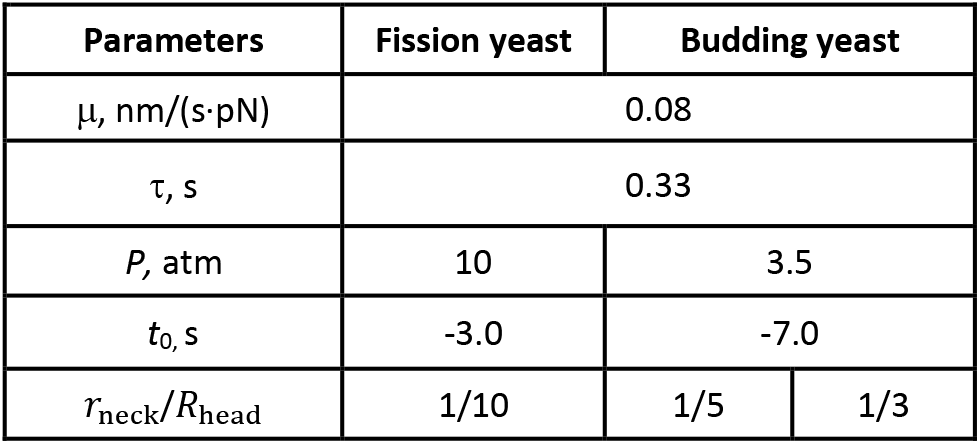
Optimized parameter sets for simulating invagination motility in fission and budding yeast.

invagination neck, is lower in budding yeast. So, with other factors being similar, the invaginations in budding yeast would likely have larger ratios *r*_neck_/*R*_head_. We solved the model for budding yeast with two values of ^*r*^_neck_/R_head_. Simulations with *r*_neck_/ R_head_ = ⅕ (*r*_neck_ = 6 nm) yielded elongation rates closer to the observations, whereas simulations with *r*_neck_/ *R*_head_ = ⅓ (*r*_neck_ =10 nm) better approximated the observed displacements (Table 4).

**Table 4.**
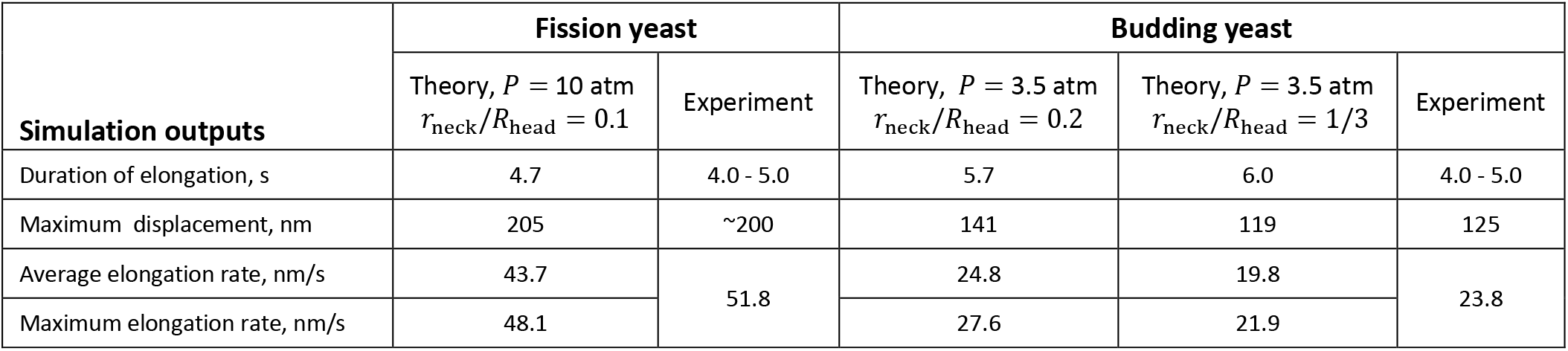
Comparison of simulation outcomes of models with optimized parameter sets with experimental data of Sun et al. (Sun et al., 2019)

| Simulation outputs | Fission yeast |  | Budding yeast |  |  |
| --- | --- | --- | --- | --- | --- |
| | Theory, $P = 10$ atm<br>$r_{\text{neck}}/R_{\text{head}} = 0.1$ | Experiment | Theory, $P = 3.5$ atm<br>$r_{\text{neck}}/R_{\text{head}} = 0.2$ | Theory, $P = 3.5$ atm<br>$r_{\text{neck}}/R_{\text{head}} = 1/3$ | Experiment |
| Duration of elongation, s | 4.7 | 4.0 - 5.0 | 5.7 | 6.0 | 4.0 - 5.0 |
| Maximum displacement, nm | 205 | ~200 | 141 | 119 | 125 |
| Average elongation rate, nm/s | 43.7 | 51.8 | 24.8 | 19.8 | 23.8 |
| Maximum elongation rate, nm/s | 48.1 |  | 27.6 | 21.9 |  |

We introduced additional minor changes to simulations run with optimized parameters, aimed at improving the overall consistency of the model and the agreement of the simulation results with experimental observations. We replaced Eq (2) with a more accurate version of *f*_resist_(*t*), that after the invagination changes shape does not drop to zero but rather approaches 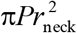,

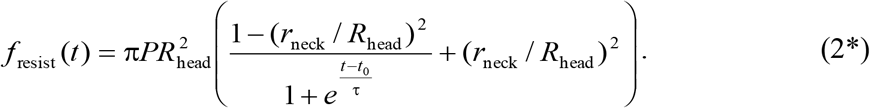

In determining durations of elongation and average elongation rates, we took into account that immediately after *t*_begin_, the invaginations elongate much slower than during the ensuing near-linear increase in displacement (see Figures 1D, 2C). Similar to the experimental study (Sun et al., 2019), we evaluated the elongation durations and rates for the fast near-linear increase of displacements, starting at the inflection point and ending when polymerized actin reaches its maximum. The inflection points of the displacement time dependencies for the budding yeast invaginations were −6.9 s with *r*_neck_/*R*_head_ = ⅓ and −6.8 s with *r*_neck_ / *R*_head_ = ⅕, and −2.8 s for fission yeast with *r*_neck_/*R*_head_ = 0.1. The corresponding end times were −1.2 s, −0.8 s, and 1.9 s.

Figure 4 compares results for budding and fission yeast obtained with the optimized parameter sets. Graphs in panels A and B illustrate how turgor pressure affects *f*_drive_(*t*) and *f*_net_(*t*) = *f*_drive_(*t*) – *f*_resist_(*t*). In panel A the dashed curves are *f*resist(*t*) described by Eq (2*) with the maximum resistance 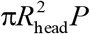 controlled by turgor pressure and thus is lower in budding yeast.

Consequently, in budding yeast elongation begins earlier during patch assembly and requires a weaker driving force, yielding lower *f*_net_ (panel B), slower elongation, and shorter displacements (panel C).

Simulation results obtained with optimized parameter sets agree reasonably well with the experimental data of Sun et al. (Table 4). The comparison reinforces the conclusion that different turgor pressures in the two yeast species are a major determinant in the observed differences of elongation rates and depths of their endocytic invaginations.

### 3. The model predicts that the peak numbers of actin are lower in budding yeast than fission yeast

Feedback between kinematics of the invagination and the accumulation of actin around the invagination explains why simulations of the model produce more polymerized actin in patches assembling under higher turgor pressures (Table 2, Figure 4D) in spite of using a fixed set of kinetic parameters for actin nucleation, polymerization, and severing (Berro et al., 2010; Nickaeen et al., 2019). Indeed, the amounts of actin polymerizing in patches under different turgor pressures are similar until the invaginations begin to move (Figure 4D), when the rate of polymerization and elongation increase abruptly (Figure 2B, 2D).

The velocities of actin filaments (and of active Arp2/3) in the immediate vicinity of a stationary initial invagination are close to zero (due to the no-slip condition reflecting the binding of actin filaments to the coat proteins), so actin dendritic nucleation is confined to a limited space around the invagination neck (Figure 2A, left), and as the density of actin filaments increases over time, polymerization slows due to excluded volume effects. Once the invagination starts to move, its elongation speed and, consequently, the velocities of the filament network in the vicinity of the invagination increase sharply, as evident from the time dependences of net driving force (Figure 4B) (recall that according to Eq (1), *u*(*t*) ∞ *f*_net_(*t*)). As a result, the space where new filaments can nucleate and grow expands as well. Therefore, the rate with which the invagination elongates influences the rate of actin accumulation in the patch, which is cumulatively reflected in the peak amounts of polymerized actin (Table 2).

The peak amounts of actin in endocytic patches estimated with traces of GFP-actin are ~ 4100 - 7500 subunits for fission yeast (Sirotkin et al., 2010; Arasada and Pollard, 2011) and ~ 3600 subunits for budding yeast (Manenschijn et al., 2019). The maximum numbers of actin in patches are higher in the simulations than the experimental values for both yeasts, but the ratios are roughly similar: the optimized fission yeast parameters yielded the maximum of 9829 polymerized subunits, whereas simulations of budding yeast produced maxima of 5637 subunits with *r*_neck_/*R*_head_ = ⅕ and 5165 subunits with *r*_neck_/*R*_head_ = ⅓. We obtained these numbers from simulation results by integrating actin densities inside an elongating cylindrical surface that encompasses the high-density filament meshwork of the patch. The white rectangular box in Figure 5 represents a cross-section of this surface.

**Figure 5.**
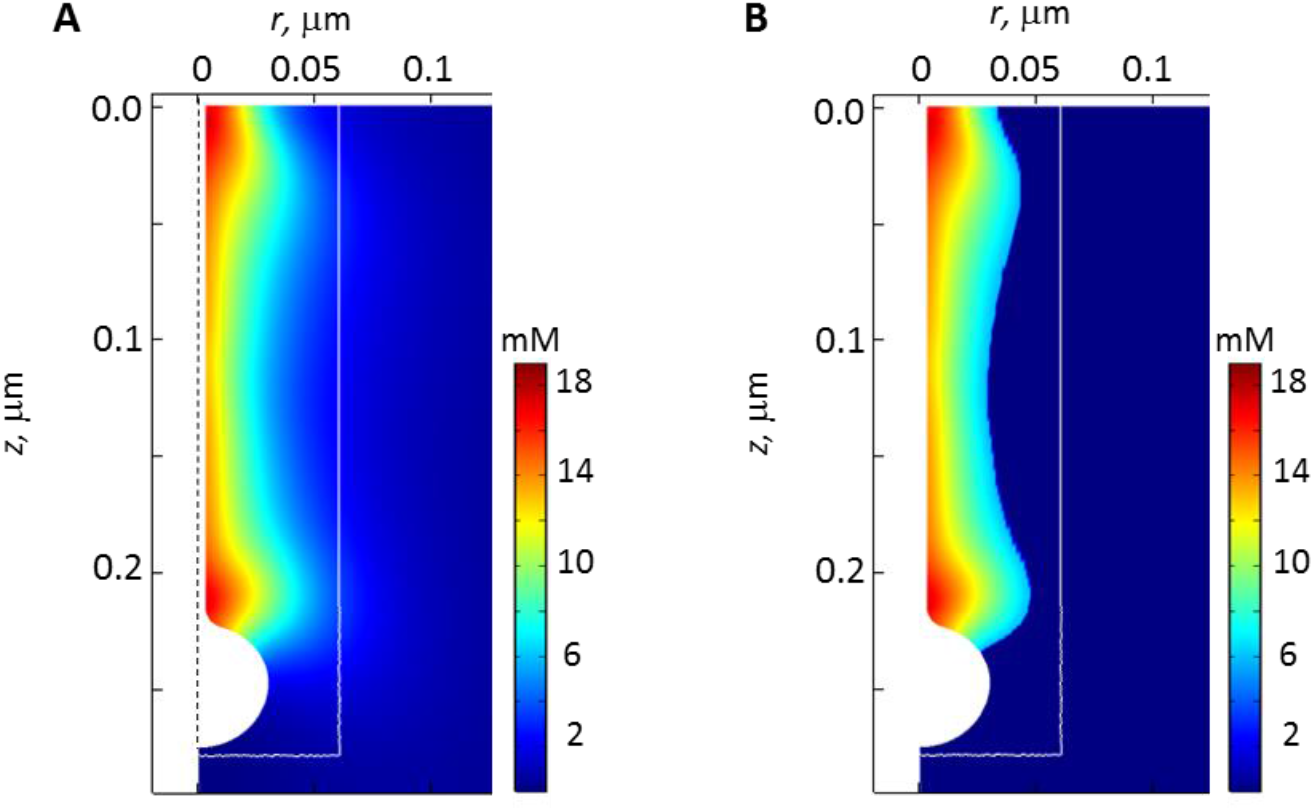
Deriving the number of actin subunits in patches from simulated actin densities. We determined the number of actin subunits in a patch by integrating actin densities within a cylinder outlined in white, which elongates with the invagination (see legend of Figure 2). (A) Snapshot of actin densities (pseudo-colors) at the time of peak actin (*t* = 1.9 s), obtained with optimized parameters for fission yeast (Table 3). (B) Actin densities from (A) exceeding 6 mM: thresholding eliminates contributions from light-blue subspaces inside the box that may not be detected experimentally.

Figure 5A is a snapshot of the actin density distribution at the time maximum patch actin in fission yeast (*t* = 1.9 s), obtained with optimized parameters (Table 3). Note that the white box includes light-blue spaces with relatively low densities of actin filaments, which may not be detected experimentally. Contributions from such spaces can be eliminated by imposing a density threshold. Figure 5B shows a patch of filament meshwork with densities exceeding 6 mM. Such thresholding lowers the number of subunits from 9829 to 5469 subunits, in the range of the experimental estimates for fission yeast. Importantly, applying the same threshold to simulation results obtained with the budding yeast parameters (Table 3) yielded a near proportionate decrease of the patch subunit count, from 5637 to 2591 subunits for *r*_neck_/*R*_head_ = ⅕ and from 5165 to 2489 subunits for *r*_neck_/*R*_head_ = ⅓, thus maintaining the approximately two-fold ratio of the numbers of subunits in the patches of the two yeasts.

The differences between the experimental estimates and simulated numbers of polymerized subunits may also reflect a modest bias against the incorporation of GFP-actin into filaments.

## Discussion

Sun et al. recorded endocytic events in budding and fission yeast in the same microscope field. The side-by-side comparison revealed remarkable similarities of the abundance of proteins at sites of endocytosis in the two yeasts, yet endocytic invaginations elongated twice as fast to twofold greater depths in fission yeast than budding yeast (Sun et al., 2019). In this study, we show that a molecularly explicit model of forces exerted by actin filaments on endocytic invaginations (Nickaeen et al., 2019) explains these differences. Counterintuitively, higher turgor pressures favor faster, deeper membrane invaginations using the same parameters of actin nucleation and polymerization.

At the core of our theory is an observation that polymerization of a dense network of actin filaments around an endocytic invagination not only produces a driving force parallel to the axis of invagination but also exerts orthogonal forces that compress the base of the invagination and stretch its middle. If sufficiently strong, these forces deform the nascent, cylindrical invagination into a flask-like shape as observed in electron micrographs of invaginations in budding yeast (Kukulski et al., 2012; Buser and Drubin, 2013). The estimates of orthogonal forces exerted by actin at the invagination base (Nickaeen et al., 2019) amount to an extra squeezing pressure with the maximum of ~ 8.5 atm, transforming the invagination shape from cylindrical to flask-like. Interestingly, the flask shapes, also predicted by the models of a membrane pulled under high turgor pressure by a point force at its tip (Dmitrieff and Nédélec, 2015; Ma and Berro, 2021), had wider necks than in electron micrographs. The orthogonal forces exerted by actin at the base of invagination could help narrow the neck.

The shape change, in turn, affects the driving and resistive forces. Since turgor pressure is isotropic, its resistance declines during the transition to a head-neck shape (Nickaeen et al., 2019). We find that this transition also changes composition of the driving force in terms of viscous and active components, due to active stresses that the polymerization of the actin filament network produces on the upper side of the invagination head. Once the driving force overcomes the declining resistive force and the invagination begins to elongate, the active component of the driving force becomes dominant and produces faster elongations and deeper invaginations (Figure 3).

Further, we find that higher turgor pressure favors faster elongations and longer displacements. For higher turgor pressure, the driving force has a chance to grow before it matches higher initial resistance. This results in higher net driving force during elongation, as the resistive force drops. Thus, the difference of turgor pressure in the two yeasts is a plausible explanation for the observed differences in motility characteristics of their endocytic invaginations.

The model also predicts, in qualitative agreement with experimental data on actin accumulation in endocytic patches (Sirotkin et al., 2010; Arasada and Pollard, 2011; Manenschijn et al., 2019), that the peak amounts of actin in the patches must be lower in budding yeast than fission yeast, because of feedback between the elongation rate and the rate of actin accumulation. Another prediction is that the ratio *r*_neck_ / *R*_head_ of the invaginations is lower in budding yeast than fission yeast.

The modeling in this study employs certain simplifications. We describe actin filament meshwork continuously in terms of concentrations of proteins participating in actin assembly, without resolving individual filaments. Given the large numbers of polymerized subunits in the patch, this approach yields reasonably accurate results, while avoiding logistical burdens of discrete stochastic simulations. In this approximation, the concentration of barbed ends serves as an estimate for the local filament density, and the ratio of the density of polymerized subunits to barbed ends estimates the local average number of subunits per filament.

Our model does not include membrane mechanics, and, therefore, we do not solve for shape dynamics. We model elongation of a head-neck invagination by increasing the length of its neck at a rate governed by Eq (1), without changing the predefined radii of the head or neck. Parameters associated with the shape of the invagination, such as the ratio of radii *r*_neck_/*R*_head_ and the time *t*_0_ around which shape change takes place, are constrained by available experimental data. In a more rigorous approach including the mechanics of the invaginated membrane (Zhang et al., 2015; Dmitrieff and Nédélec, 2015; Ma and Berro, 2021), the solution of the model would yield the dynamic geometry of the invagination along with the distributions of velocities and densities of polymerized actin. Formulating such a model will be possible once detailed knowledge is available on the composition and rheological properties of the endocytic membrane and its protein coat.

Even with its simplifications, our model reproduced experimentally observed elongation rates and displacements of endocytic invaginations in fission and budding yeast, and uncovered the connection between turgor pressure and invagination motility.

## Acknowledgements

Research reported in this publication was supported by National Institute of General Medical Sciences of the National Institutes of Health (NIH) under award numbers R01GM026338, P41GM103313 and R01GM115636, and by the National Science Foundation (NSF) under award number MCB171605. The content is solely the responsibility of the authors and does not necessarily represent the official views of the NIH or the NSF. B.M.S. thanks Leslie Loew for continuing support. The research presented in this paper was supported by the systems, services, and capabilities provided by the University of Connecticut High Performance Computing (HPC) facility.

## Supplemental Text

### Model and results: mathematical details

#### 1. Nucleation and polymerization kinetics and transport of actin

The model combines detailed kinetics of actin nucleation and polymerization (Berro et al., 2010) and mechanics of actin filament meshwork approximated as a visco-active gel (Kruse et al., 2005). It is formulated in a continuous approximation that does not resolve individual filaments, but rather describes a distribution of actin in the patch by a continuous density of polymerized subunits ρ as a function of spatial location **r** and time *t*, ρ(**r**, *t*). The actin density p is determined by concentrations of all of the species of actin in an actin patch including the newly polymerized ATP-bound subunits (FATP), the subunits aged by ATP hydrolysis and phosphate dissociation (FADP), the subunits bound by cofilin (FCOF), the filaments barbed-ends, both active and capped (BEa and BEc, respectively), and the slowly depolymerizing pointed ends (PE): ρ = *n_A_*∑_*X*_ [*X*], where *X* stands for FATP, FADP, FCOF, BEa, BEc, and PE; [*X*] is the concentration of molecule *X* in μM; the prefactor *n_A_* converts the concentration in μM into the density expressed in molecules per μm^3^ (*n_A_* = 602 μm^−3^/μM).

The concentrations [*X*] are governed by transport-reaction equations of the following type,

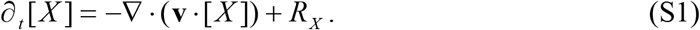

The first term in Eq (S1) is the rate of change of [*X*] due to advective flow of actin meshwork characterized by the vector field of local time-dependent velocities **v** = **v**(**r**, *t*). Like [*X*] and surface densities [*Y*] of actin-nucleating species (see below), actin velocities **v**(**r**, *t*) are the model variables; they are governed by the mechanics of actin filament meshwork described in the next section. The second term in Eq (S1) is the sum of rates of all reactions affecting a given *X*. The wiring schematic of reactions involved in nucleation, polymerization, and severing of actin, and capping of the barbed ends of actin filaments is shown in Figure S1. The diagram includes two more volume species that do not directly contribute to ρ(**r**, *t*): the active Arp2/3 complex (ActiveArp) and the Arp2/3 complex in the filamentous actin network (FArp). While the equation governing FArp is of the form of Eq (S1), the equation for ActiveArp has an additional diffusion term:

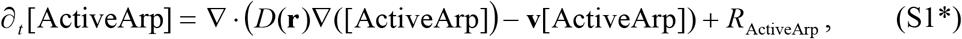

where *D***(r)** is nonzero only in the vicinity of the rings of nucleation promoting factors (NPFs), as described in the next section (for reasoning and details, see (Nickaeen et al., 2019)).

Reaction steps leading to formation of ActiveArp occur on the surface of the membrane, where they localize to the rings occupied by the nucleation promoting factors WASp (Figure S1). They involve dimers of WASp bound to G-actin monomers (WGD), Arp2/3 ternary complexes consisting of Arp2/3 complex bound to WGD (ArpTernCompl), and activated Arp2/3 ternary complexes (FArpTernCompl), whose surface densities are described by rate equations, *δ_t_*[*Y*] = *R_Y_*, where [*Y*] is the surface density of a membrane-bound protein *Y* in molecules per squared micron. Note that while these variables are governed by ordinary differential equations, they also depend on spatial coordinates, given that *R_Y_* are nonzero at the rings occupied by WASp, and *R*_ArpTernCompl_ depends on [FATP] and [FADP] at the plasma membrane.

**Figure S1.**
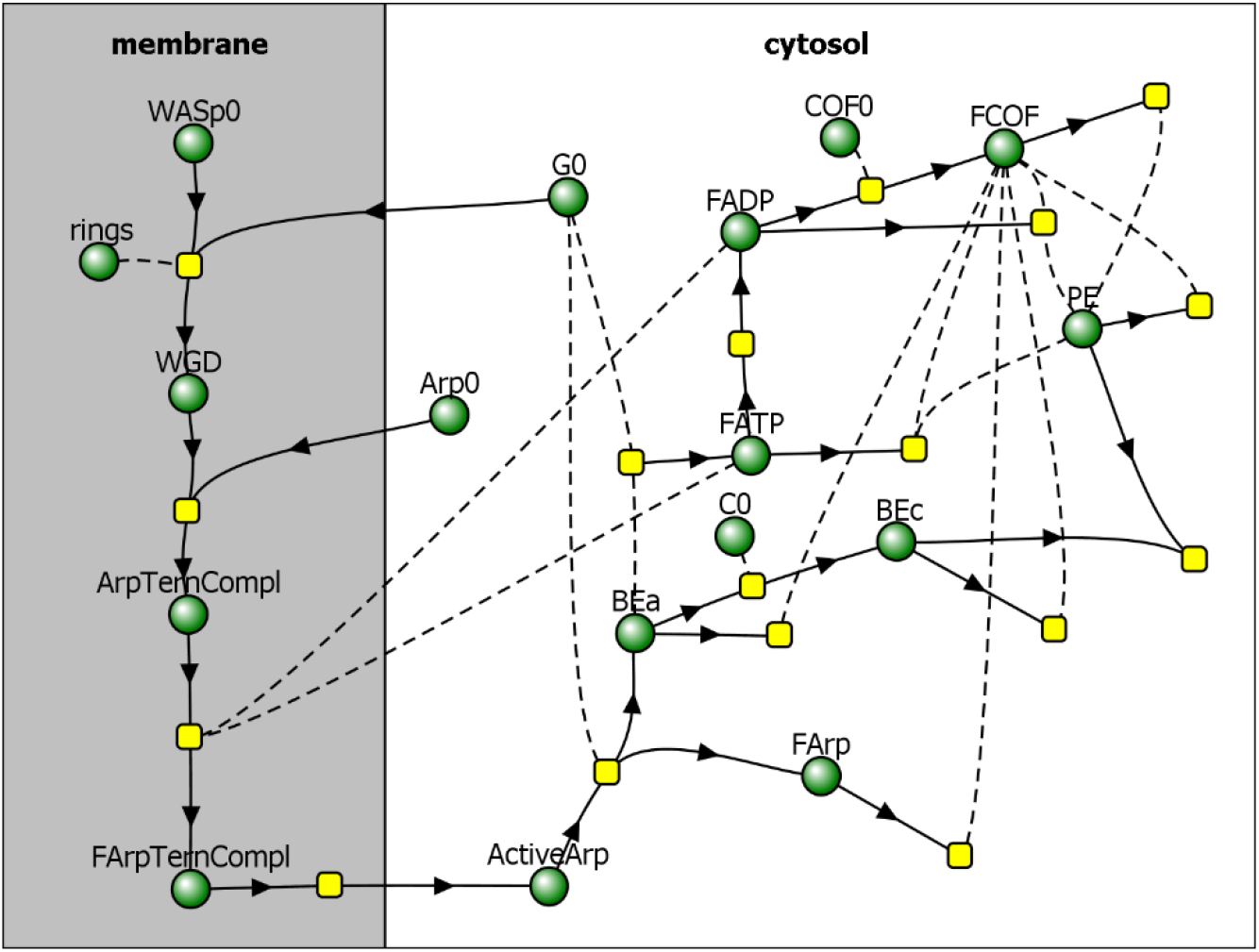
Reaction diagram of nucleation and polymerization of patch actin proposed in (Berro et al., 2010), with added partitioning of species between membrane and cytosol (adapted from (Nickaeen et al., 2019)). Directions of arrows towards or from reaction nodes (yellow squares) determine roles of molecular species (green circles) in a particular reaction as reactants or products, and reactions without products describe disappearance of reactants from the patch. Species connected to reactions by dashed curves act as ‘catalysts’, i.e. they are not consumed in those reactions.

Reaction rates structure in *R_X_* and *R_Y_* are as in (Berro et al., 2010). The on (+) and off (−) rate constants of polymerization, capping, cofilin binding, and cofilin-dependent severing from (Berro et al., 2010) were modified by the factor α(**r**, *t*) = (1 – ρ(**r**, *t*)/ρ_max_)^½^, to reflect the effect of molecular crowding; and the rates of polymerization and capping were modified by an additional factor β(**r**, *t*) = 1 – ρ(**r**, *t*)/ρ_max_, to take into account that polymerization and capping slow down under load, where ρ_max_ = (4πδ^3^/3)^−1^ (δ is the length of actin subunit; for derivation details, see (Nickaeen et al., 2019)). The expressions of *R_X_* and *R_Y_* used in computations are shown below:

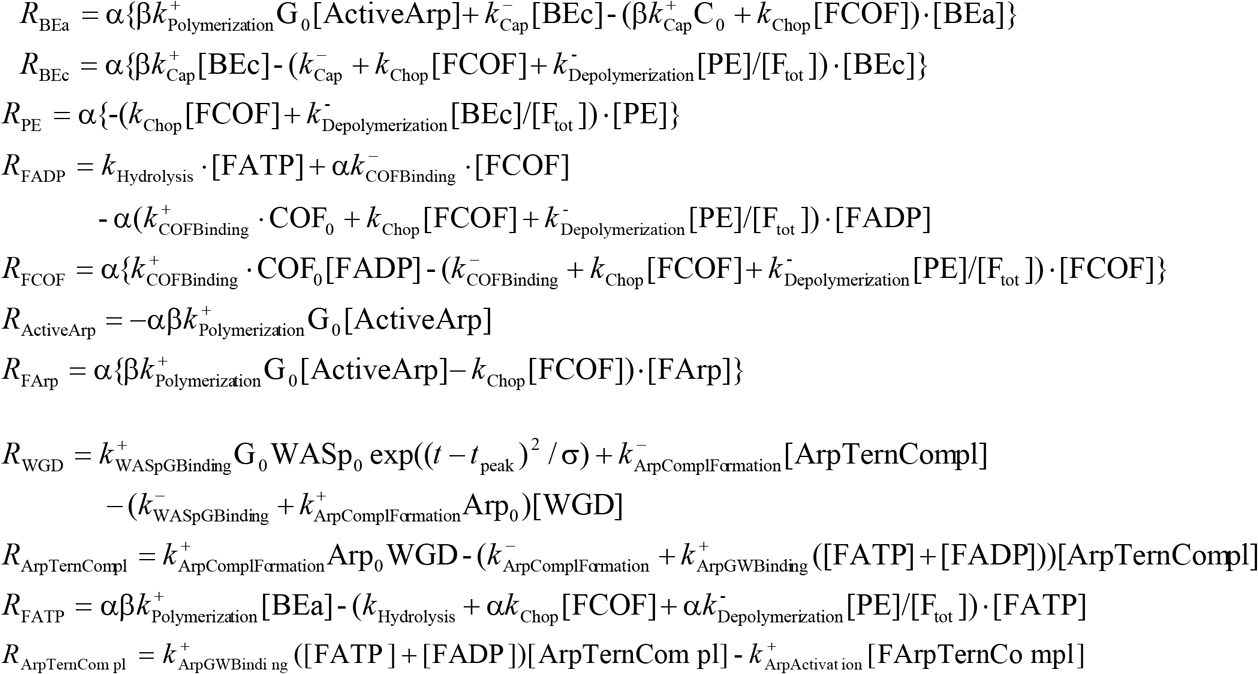

In the expressions above, F_tot_ = ρ / *n_A_* = [FATP] +[FADP] +[FCOF] +[BEa] +[BEc] +[PE] and the zero subscript denotes constant concentrations and surface density. Table S1 summarizes the values of constants used in the reaction rate expressions and initial conditions for Eqs (1) and for the rate equations describing actin nucleation. For their sources, see Tables 1 and 2 of (Berro et al., 2010). Table S1 also includes the constants involved in actin mechanics: the actin subunit length δ and the constants κ_active_ and κ_viscous_ setting the scales of active and viscous stress and of actin velocities; for derivation of κ_active_ and κ_viscous_, see (Nickaeen et al., 2019).

#### 2. Mechanics of actin meshwork

We interpret mechanics of actin meshwork as that of a compressible visco-active fluid (Kruse et al., 2005; Prost et al., 2015). In a viscosity-dominated environment of the actin patch, actin velocities **v**(**r**, *t*) are governed by the balance of local active and viscous forces (per unit volume) **f**_active_ and **f**_viscous_ (Kruse et al, 2005):

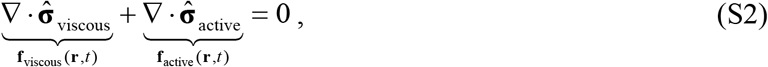

where 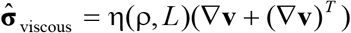 and 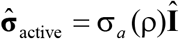 are the viscous and active stress tensors, respectively (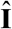 is the unit tensor). In the expression for the viscous stress tensor, (∇**v**)^*T*^ is the transpose of the velocity gradient tensor ∇**v**, and *L* = *L*(**r**,*t*) is the local average length of actin filaments, computed as *N*δ, with the number of subunits in a filament *N* = *N*(**r**,*t*) = [Ft_tot_]/([BEa] +[BEc]); η(ρ,*L*) is the dynamic viscosity of the actin filament meshwork, and σ_*a*_(ρ) is the energy (per unit volume) stored in the meshwork during polymerization. By analyzing rheological properties of overlapping actin filaments (Gardel et al., 2003; Kasza et al., 2010; Tseng and Wirtz, 2004; Mullins et al., 1998), we derived the following constitutive relations for η(ρ,*L*) and σ_*a*_(ρ):

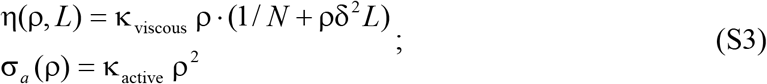

for derivation details, see (Nickaeen et al., 2019).

**Table S1.** Model parameters

| Parameter | Value | Units |
| --- | --- | --- |
| $k_{\text{ArpActivation}}^+$ | 0.5 | $\text{s}^{-1}$ |
| $k_{\text{ArpComplFormation}}^+$ | 0.8 | $\mu\text{M}^{-1}\text{s}^{-1}$ |
| $k_{\text{ArpComplFormation}}^-$ | 0.74 | $\text{s}^{-1}$ |
| $k_{\text{ArpGWBinding}}^+$ | 0.3 | $\mu\text{M}^{-1}\text{s}^{-1}$ |
| $k_{\text{ArpGWBinding}}^-$ | 0.001 | $\text{s}^{-1}$ |
| $k_{\text{Cap}}^+$ | 7 | $\mu\text{M}^{-1}\text{s}^{-1}$ |
| $k_{\text{Cap}}^-$ | 0.004 | $\text{s}^{-1}$ |
| $k_{\text{COFBinding}}^+$ | 0.0085 | $\mu\text{M}^{-1}\text{s}^{-1}$ |
| $k_{\text{COFBinding}}^-$ | 0.005 | $\text{s}^{-1}$ |
| $k_{\text{Polymerization}}^+$ | 11.6 | $\mu\text{M}^{-1}\text{s}^{-1}$ |
| $k_{\text{Depolymerization}}^-$ | 0.25 | $\text{s}^{-1}$ |
| $k_{\text{WASpGWBinding}}^+$ | 42.9 | $\mu\text{M}^{-1}\text{s}^{-1}$ |
| $k_{\text{WASpGWBinding}}^-$ | 25.7 | $\text{s}^{-1}$ |
| $k_{\text{Chop}}$ | 0.003 | $\mu\text{M}^{-1}\text{s}^{-1}$ |
| $k_{\text{Hydrolysis}}$ | 0.3 | $\text{s}^{-1}$ |
| $\text{Arp}_0$ | 1.3 | $\mu\text{M}$ |
| $\text{C}_0$ | 0.8 | $\mu\text{M}$ |
| $\text{COF}_0$ | 40 | $\mu\text{M}$ |
| $\text{G}_0$ | 21.6 | $\mu\text{M}$ |
| $\text{WASp}_0$ | 5192* | molecules/$\mu\text{m}^2$ |
| $t_{\text{peak NPF}}$ | 17.2** | s |
| $\sigma$ | 33.5 | $\text{s}^2$ |
| $\kappa_{\text{active}}$ | $3.69 \times 10^{-3} n_A^{-2}$ | $\text{Pa}/(\mu\text{M})^2$ |
| $\kappa_{\text{viscous}}$ | $3.93 n_A^{-2}$ | $\text{Pa} \cdot \text{s}/\mu\text{M}$ |
| $\delta$ | 2.7 | nm |

\* Maximum surface density of free WASp in the rings is the number of WASp molecules, that corresponds to concentration 0.23 $\mu\text{M}$ in a sphere of radius 0.15 $\mu\text{m}$ (Berro et al., 2010), divided by area of two rings, $4\pi r_{\text{neck}} w$ , where $w$ is the width of one ring. The value is shown for $w = 5$ nm, used in all simulations, and $r_{\text{neck}} = 3$ nm (some simulations were run with neck radii of 6 and 10 nm).
\*\* Upon offsetting by -20 s, $t_{\text{peak NPF}}$ becomes -2.8 s (Berro et al., 2010).

#### 3. Computation domain, boundary and initial conditions

Equations (S1), (S1*), and (S2) are to be solved in a section of the cell embedding a head-neck invagination, see the diagram of a 3D computational domain Ω in Figure S2A (drawn not to scale). The domain is bounded by the plasma membrane δΩ_membrane_, which includes the invagination, and two additional surfaces that define its vertical cylindrical sides and the horizontal floor. In order for the conditions at the outer limits of the domain, which are arbitrary, to not have significant effect on the results, we solved the model in a sufficiently large, 0.5 μm in each coordinate direction, neighborhood of the invagination. Note that because the invaginated portion of the domain boundary can move, the shape of the computational domain generally changes with time.

Because the model geometry, localization of membrane-bound species, and corresponding fluxes remain axisymmetric throughout the elongation process, solutions of the model are also axisymmetric. This reduces the problem to solving an equivalent 2D model in cylindrical coordinates (*r*,*z*) in the domain Ω depicted Figure S2B (not to scale). The full 3D geometry is then restored for any time by revolving the 2D domain Ω around the axis of symmetry *r* = 0.

The equations (S1) for all volume variables, except for [ActiveArp], are of the hyperbolic type and subject to zero-flux boundary conditions at ∂Ω_cell_membr_ ∪ ∂Ω_invagination_ ∪ ∂Ω_symm_axis_ (Figure S2B). For implementation details, see Supplemental Text of (Nickaeen et al., 2019).

**Figure S2.**
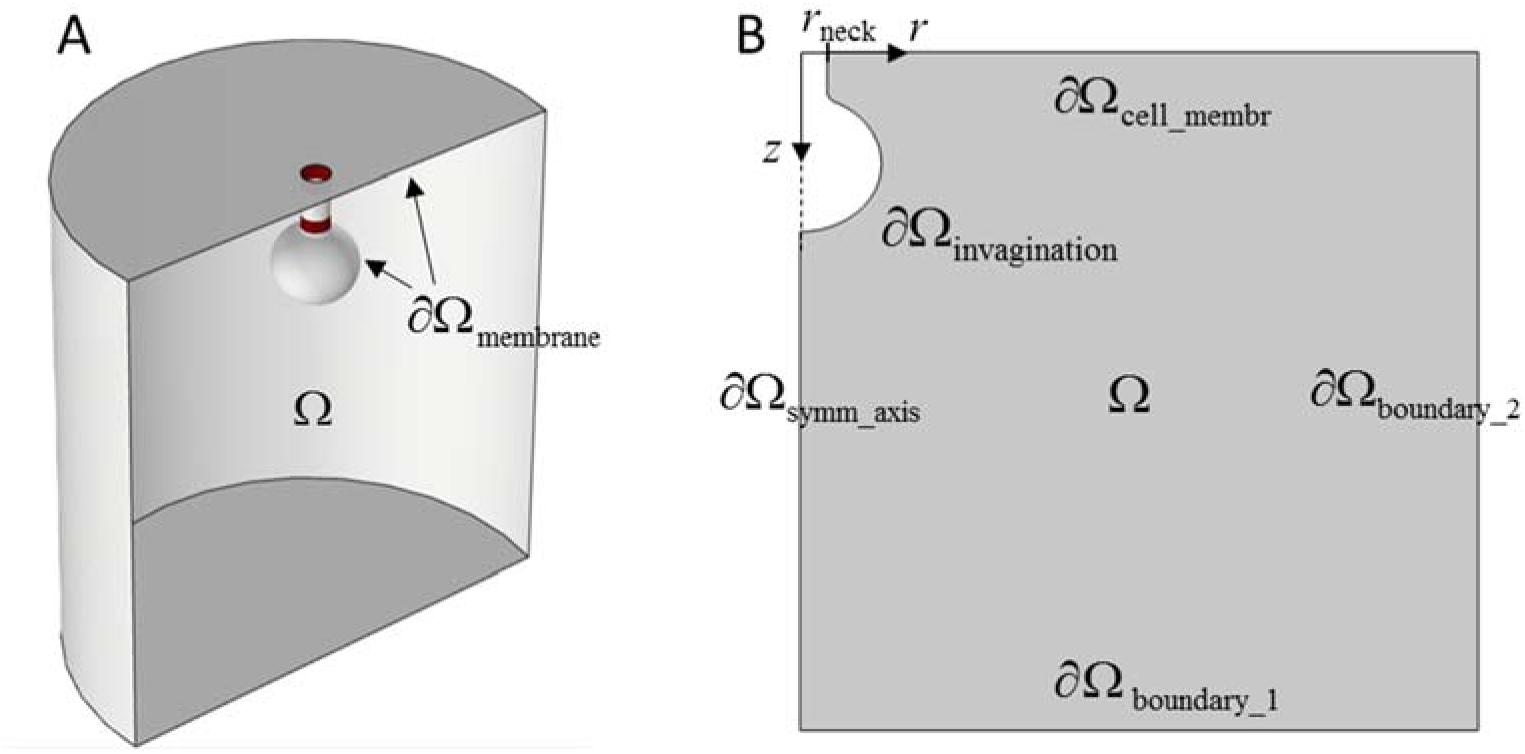
Computational domain. (A) 3D diagram of the fragments of cell, Ω, and plasma membrane, including invagination, ∂Ω_membrane_, comprising computational domain (not to scale). Two rings of NPFs are shown in dark red. When invagination elongates, both ∂Ω_membrane_ and Ω change with time. (B) Computational domain of the equivalent 2D problem in (*r,z*) coordinates (not to scale). The domain boundary ∂Ω includes: fragment of cell membrane ∂Ω_cell_membr_, invagination ∂Ω_invagination_, fragment of axis of symmetry, ∂Ω_symm_axis_, and horizontal (∂Ω_boundary_1_) and vertical (∂Ω_boundary_2_) outer boundaries; ∂Ω_membrane_ = ∂Ω_cell_membr_ ∪ ∂Ω_invagination_.

Eq (S1*), governing [ActiveArp], is of the parabolic type, because it includes a diffusion term. Therefore, boundary conditions for this equation must be specified at all boundaries in Figure S2B. At ∂Ω_invagination_, it must satisfy the Rankine-Hugoniot boundary condition (Novak and Slepchenko, 2014),

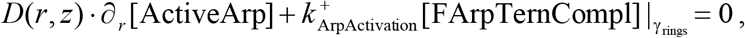

where γ_rings_ are the locations on ∂Ω_invagination_ occupied by the rings of the NPFs. Recall that in the equation above, [ActiveArp] is measured in μM, whereas [FArpTernCompl] is in molecules per squared micron. Note also the disappearance of the advection term from the boundary condition, which is due to the no-slip boundary condition for Eq (S2) that we describe later in this section.

The diffusion coefficient of [ActiveArp] *D*(*r*,*z*) is defined as follows:

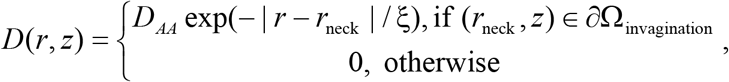

with *D_AA_* = 0.001 μm^2^/s and = ξ nm. The value of ξ is on the order of mesh sizes used in the computations; this parameter has little effect on the solution, because ActiveArp quickly converts to BEa. Because *D*(*r*,*z*) is nonzero only in the tight space near the invagination, varying *D_AA_* by several orders of magnitude did not change the simulation outcome in any significant way.

The boundary conditions for [ActiveArp] at ∂Ω_cell_membr_ ∪ ∂Ω_symm_axis_ were zero-flux. At the remaining two boundaries, ∂Ω_boundary_1_ and ∂Ω_sboundary_2_, we applied outflow boundary conditions. For reasons explained in Supplemental Text of (Nickaeen et al., 2019), we specified the outflow boundary conditions at ∂Ω_boundary_1_ ∪ ∂Ω_boundary_2_ for all volume variables.

Similar to the Stokes equation for a Newtonian fluid, Eq (S2) is elliptic in nature and hence requires that boundary conditions be specified at all boundaries of Ω. At ∂Ω_cell_membr_ and ∂Ω_invagination_, we applied the no-slip boundary condition,

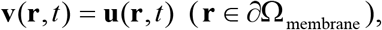

where the membrane velocity **u**(**r**,*t*) may be nonzero only for the points of ∂Ω_invagination_. In the model, these points move with the same velocity, **u**(*t*) = (*u_r_*(*t*), *u_z_*(*t*)) = (0, *u*(*t*)), where *u*(*t*) is governed by Eq (1) of the main text. At ∂Ω_symm_axis_, velocities **v**(**r**,*t*) were set to zero, and at ∂Ω_boundary_1_ ∪ ∂Ω_boundary_2_, the velocities satisfied the zero-stress boundary condition: 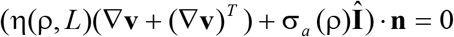

The initial actin velocities and concentrations of all molecular species, except for [FADP], [BEa], and [PE], were set to zero. Small initial values assigned to [FADP], [BEa], and [PE] reflected a small number of seed filaments (Chen and Pollard, 2013).

#### 4. Computation of the driving force

The elongation rate is determined in our model by the interplay of the driving force *f*_drive_(*t*) produced by the stresses in the assembling actin patch, and the resistive force *f*_resist_ (*t*) due to turgor pressure (the latter is defined by Eqs (2) and (2*) of the main text). From fluid mechanics, the driving force is found by integrating the viscous and active tress tensors projected on the outward normal (Landau and Lifshitz, 1989),

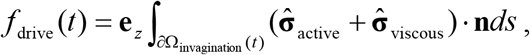

where the integral is carried over the time-dependent invaginated membrane ∂Ω_invaginaton_(*t*), **n** = (*n_r_*,*n_z_*)^*T*^ is the outward unit normal vector to the membrane (directed towards the interior of Ω), **e**_*z*_ is the unit vector parallel to the axis of symmetry, and *ds* is the area of an infenitesimal element of the invaginated membrane.

#### 5. Numerical solution of the model

We solved the equivalent 2D model numerically using a moving-mesh solver of COMSOL® Multiphysics (COMSOL Multiphysics, 2015). The solver utilizes Arbitrary Lagrangian Eulerian (ALE) methods based on finite-element (FE) discretization. The ALE methods are described in numerous publications, see e.g. (Donea et al., 2004).

In an ALE simulation, the computational grid points move with the velocities defined as follows. At the moving interfaces, they coincide with the velocities of the points of the interface. In the interior of the domain, the velocities of the grid are arbitrary, so long as they comprise the smooth vector field that would maintain mesh quality throughout the simulation, while preserving mesh connectivity. Correspondingly, the governing equations formulated with respect to a fixed (Eulerian) coordinate system should be reformulated based on the ALE methodology. For the implementation of our model in the ALE framework and details of its FE formulation, see Supplemental Text of (Nickaeen et al., 2019).

#### 6. Effects of the timing of resistance reduction on beginning of elongation and maximum driving force

The timing of the resistance descent *t*_0_ affects the time *t*_begin_, at which the invagination begins to elongate, and the magnitude of the driving force at *t* = *t*_begin_. Indeed, *t*_begin_ satisfies the equaiton *f*_drive_(*t*) = *f*_resist_(*t*), so that *t*_begin_ and *f*_drivel_(*t*_begin_) are the coordinates of a point where the graphs of *f*_drive_(*t*) and *f*_resist_(*t*) intersect (Figure S3). As *t*_0_ increases, the point of intersection ‘slides’ up along the graph of *f*_drive_(*t*), yielding higher *t*_begin_ and *f*_drive_(*t*_begin_). However, the effect saturates, once the point of intersection approaches the upper plateau of *f*_resist_(*t*), as the ‘sliding’ along *f*_drive_(*t*) slows down and eventually comes to a stop. Thus, the highest *f*_drive_(*t*_begin_) is 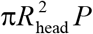 and 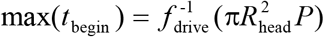, where 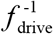 denotes the inverse function of *f*_drive_(*t*). Both limits are controlled by turgor pressure.

**Figure S3.**
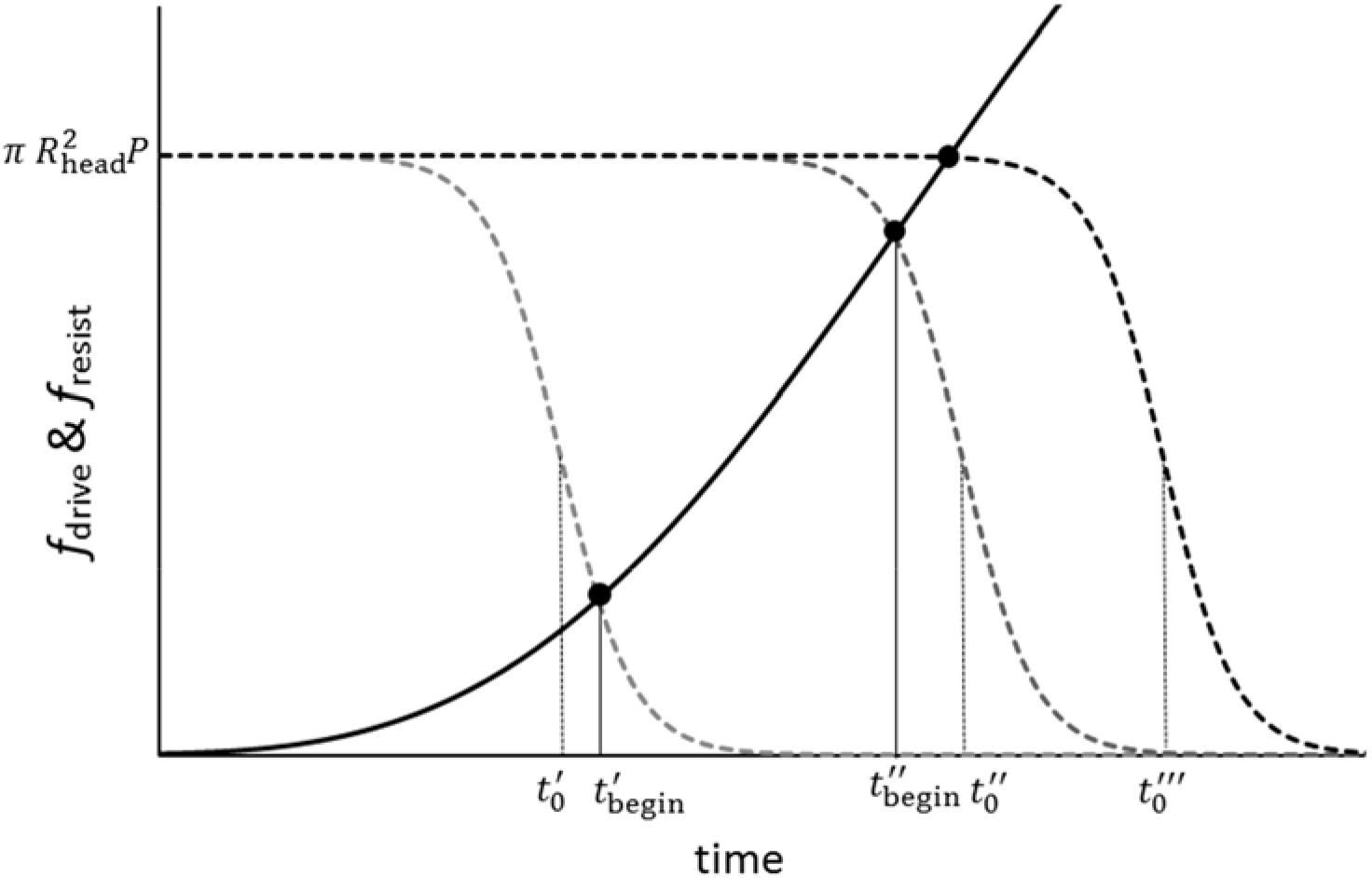
Effects of *t*_0_ on *t*_begin_ and *f*_drive_(*t*_begin_). As *t*_0_ increases, the graph of *f*_resist_(*t*) (dashed curve) shifts to the right (Eq (2) of the main text), and the coordinates of its intersection with the graph of *f*_drive_(*t*) (solid curve), *t*_begin_ and *f*_drive_(*t*_begin_), also go up. However, this effect saturates, once the point of intersection reaches the upper plateau of *f*_resist_(*t*), as is the case with 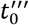; 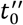 is an example of a ‘near-saturation’ value of *t*_0_.

The effect of *t*_0_ on *t*_begin_ and *f*_drive_(*t*_begin_) begins to saturate when *t*_begin_ approaches an inflection point of the rate of change of *f*_resist_(*t*), as illustrated by 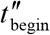 in Figure S3. From Eq (2) of the main text, the respective root *t*_*_ of δ^3^*f*_resist_(*t*) = 0 is the solution of 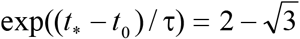, and 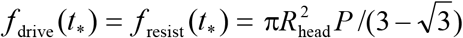. Then the ‘near-saturation’ values of *t*_0_ are 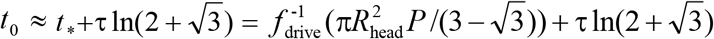.

## Footnotes

1 All simulations in our previous study were run with *μ* = 0.04 nm/(s·pN), not *μ* = 0.4 nm/(s·pN) as misstated in (Nickaeen et al., 2019). The typographical error, made through the fault of the authors, had no effect on the results of (Nickaeen et al., 2019).

